# Evaluating gene representation in spatial transcriptomics across pre-designed panels

**DOI:** 10.1101/2025.04.07.647642

**Authors:** Heewon Seo, Roman Krawetz

## Abstract

Spatial transcriptomics (ST) has revolutionized our understanding of gene expression within tissues by preserving spatial context. Over the past few years, this technology has led to a number of paradigm-shifting discoveries that have enabled a more comprehensive understanding of cellular functions and interactions in normal and diseased states. However, ST technologies still face challenges related to resolution, sensitivity, and technical variability. In this study, we evaluate the read coverage of commercialized pre-designed panels using publicly available ST datasets generated from the 10X Genomics and Nanostring platforms. We introduce the Coverage Index (CI) as a quantitative metric to assess the representation of established gene signatures across multiple ST datasets. Our findings reveal that cancer-related gene lists exhibit the highest CI values, while genes encoding for ligands and receptors tend to have low coverage. Additionally, CI analysis can help highlight intrinsic biases in gene panel design, influencing the detection capacity and thereby downstream comprehension of certain biological pathways. The insights gained from this study provide a framework for assessing ST panel performance and optimizing gene panels for future spatial transcriptomic applications.

## Introduction

Understanding the spatial organization of gene expression within tissues is essential for deciphering cellular functions, interactions, under both homeostatic and disease states. Conventional transcriptomic approaches, such as bulk and single-cell RNA-seq (scRNA-seq), provide valuable insights into gene expression but discard the spatial information which is critically essential for the understanding of complex tissues and organ systems^1^. Spatial transcriptomics (ST) has emerged as a powerful technology that bridges the gap between transcriptomic analysis and tissue architecture, enabling researchers to dissect the complex molecular landscapes of tissues^2^. However, one of the challenges remaining is the trade-off between resolution and sensitivity^3^. While ST platforms achieve near-or sub-cellular resolution, they often suffer from low transcript detection efficiency, making it difficult to capture rare or low-abundance transcripts^4,5^. Additionally, ST platforms rely on specialized instrumentation, complex sample preparation, and sequencing depth, leading to significant technical variability^6^.

Spatial transcriptomics is a powerful technology that maps gene expression within the spatial context of tissues, offering insights into cellular organization, tissue architecture, and microenvironment interactions. It has revolutionized biology and medicine by enabling the study of complex tissues, tumor microenvironments, and developmental biology, and by advancing precision medicine through biomarker discovery and targeted therapies. However, spatial transcriptomics is susceptible to biases introduced at various stages of the workflow. Changes in sample preparation, such as tissue fixation and sectioning, can alter RNA integrity and expression profiles. Human error in handling samples or during imaging and data analysis can lead to inconsistencies. Additionally, platform selection (e.g., slide-based vs. bead-based methods) and panel design (e.g., targeted vs. whole-transcriptome panels) influence data resolution and coverage, potentially introducing bias. Rigorous standardization of protocols, quality control measures, and validation strategies are crucial to minimize these biases and ensure reproducible and accurate results.

To overcome these challenges, the field has developed a number of quality control metrics that have resulted in the gradual improvement in data quality over time, yet there are still gaps that need to be addressed. One such area is technical biases, notably a consistent pattern of genes exhibiting either high or low coverage irrespective of sample/tissue type. To address this, we investigated and characterized the capabilities of pre-designed panels using publicly available ST datasets: seven 10X Genomics Xenium In Situ with the Prime 5k panel and 16 Nanostring GeoMx DSP (Digital Spatial Profiler) datasets^6–10^ using the human whole transcriptome atlas (WTA) panel. We first obtained raw count matrices and aggregated them across samples within each study. Subsequently, we compared gene ranks with established databases, i.e., MSigDB (https://www.gseamsigdb.org/), and assessed which gene lists were abundant using a binary classifier. The classifier delivers the coverage index (CI) and we finally compared CI values across various studies and databases to identify gene lists which are over/underrepresented.

## Results

### Evaluating the capacity of a pre-designed panel

For our analysis, we utilized raw count matrix profiles to calculate CI values across 23 distinct studies using ten gene databases (Fig. 1a). Higher CI values denote greater coverage among gene members within a representative gene list, while lower CI values indicate reduced coverage. A CI value of 0.5 indicates a random distribution regarding read counts. To further explore the efficacy of cell-cell interaction analysis, we compared these results with those derived from lig- and and receptor genes^11^ (Fig. 1b). It was anticipated that studies utilizing the Xenium Prime 5k would exhibit lower CI values than those obtained from GeoMx WTA, owing to the difference in the number of genes included in the relative panels. While ligand genes within the Xenium context displayed lower coverage than their receptor counterparts, GeoMx data revealed comparable CI values for both ligand and receptor gene groups. An intriguing observation was that *fresh frozen* samples did not yield greater CI values for these genes, even though they captured approximately ten-fold more transcripts per 100 µm^2^ than formalin-fixed, par-affin-embedded samples; this finding suggests that CI values may reflect inherent technical biases linked to the design of the gene panels vs. differences in sample preparation and/or handling.

**Figure 1.**
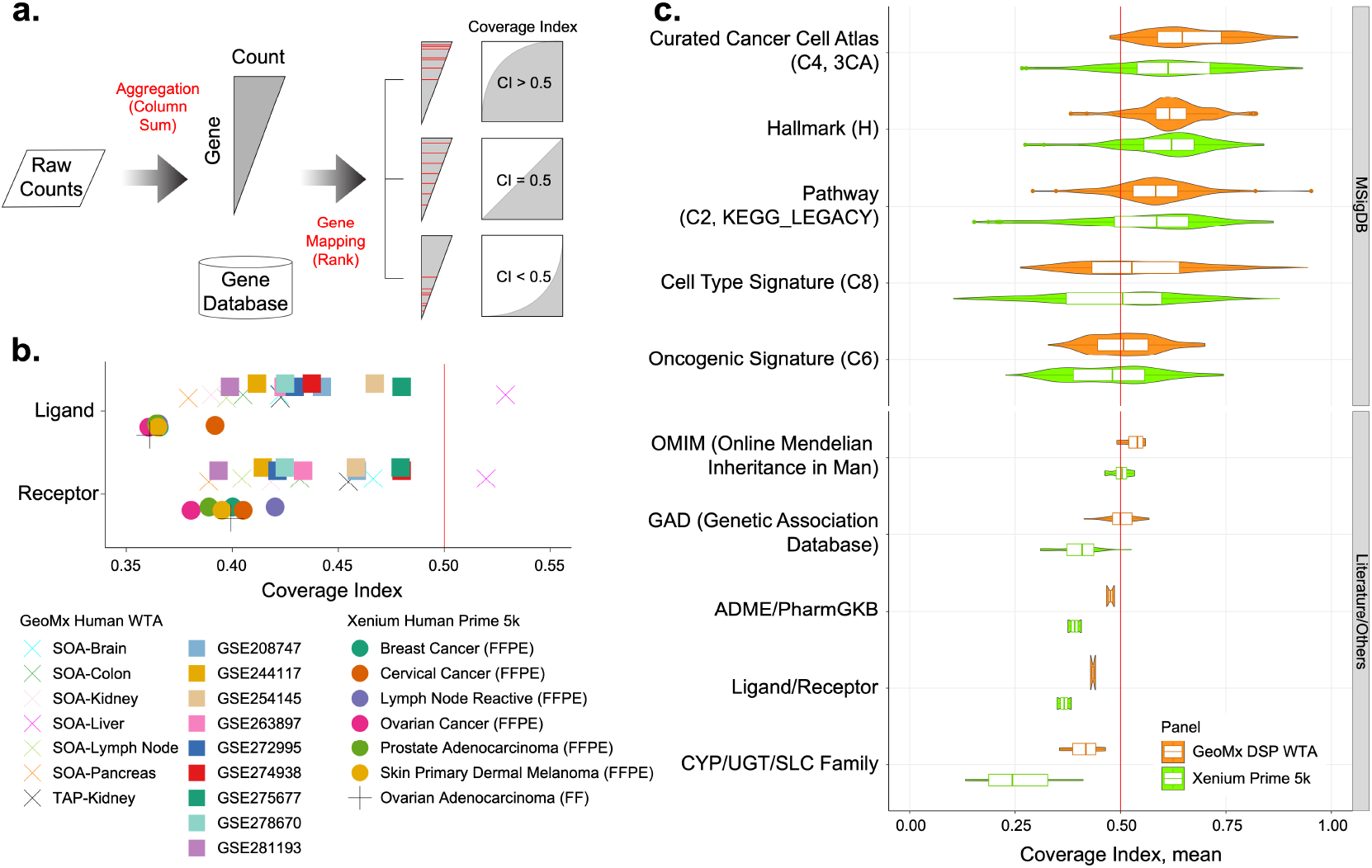
Evaluation of the pre-designed gene panel utilizing the coverage index (CI). **(a)** A schematic representation outlining the calculation of the CI. **(b)** CI values for 23 cohorts examining ligand and receptor genes. **(c)** Mean CI values across cohorts for ten different gene databases.

Furthermore, CI values for ligand/receptor genes were predominantly below 0.5, with one exception noted in the Spatial Organ Atlas (SOA)-Liver. In contrast, most can-cer-related gene lists, i.e., Cancer Cell Atlas (C4, 3CA), exhibited CI values exceeding 0.5 (Supplementary Fig. S1), suggesting that analyzing ligand/receptor genes may be more challenging due to low coverage compared to investigating cancer cell-related genes. Correlation coefficients calculated among the studies using aggregated raw counts revealed that Spearman’s rho in the Xenium panel consistently exceeded 0.75 across all comparisons, while GeoMx studies similarly demonstrated positive correlations (Supplementary Fig. S2).

### Cancer-associated genes captured a higher number of reads compared to other gene categories

We computed the mean CI values across studies to evaluate coverage in ten diverse gene databases (Fig. 1c). The Curated Cancer Cell Atlas (C4, 3CA) member genes illustrated the highest coverage across both Xenium and GeoMx platforms. At the same time, the Oncogenic Signature (C6) exhibited marginal CI values near 0.5. It is critical to note that the Curated Cancer Cell Atlas data originated from scRNA-seq, whereas the Oncogenic Signatures were derived from microarray analyses. Examining the Hallmark gene sets, we found that four of the top five gene lists— MYC_TARGETS_V1, TGF_BETA_SIGNALING, PROTEIN _SECRETION, and OXIDATIVE_PHOSPHORYLATION— were shared across both panels. Conversely, three of the bottom five gene lists also aligned: PANCREAS_BETA _CELLS, SPERMATOGENESIS, and KRAS_SIGNALING _DN, indicating that such signature genes are relatively comparable across Xenium and GeoMx panels. When analyzing the Kyoto Encyclopedia of Genes and Genomes (KEGG) pathways, distinct strengths were noted for each panel; for instance, the mean CI values for the GRAFT_VERSUS_HOST_DISEASE were recorded at 0.2969 for Xenium and 0.6130 for GeoMx (Supplementary Table S1). Conversely, the mean CI values for the GLYCOSPHINGOLIPID_BIOSYNTHESIS_GANGLIO _SERIES were 0.8622 for Xenium and 0.5585 for GeoMx. The Genetic Association Database (GAD) has cataloged genes associated with different types of diseases, revealing that cancer-related genes exhibit the highest median CI values compared to other gene categories (Supplementary Fig. S3). Conversely, the Xenium panel demonstrated a lower CI in detecting genes belonging to the CYP, UGT, and SLC protein families, primarily due to the intricate nature of paralogous relationships within these gene families. Consistent with this observation, *CYP2A6* exhibited the lowest read count among all formalin-fixed, paraffinembedded samples analyzed within the Xenium studies (Supplementary Fig. S4).

### Ligand/receptor pair analysis using the Xenium In Situ and hallmark signatures

The ligand/receptor pair analysis using Xenium is pivotal for achieving subcellular spatial resolution and understanding cell-cell interactions. To identify known ligand/receptor pair(s), CellChat v2^11^ was employed, facilitating a comparison between ligand/receptor genes and the Prime 5k genes (Supplementary Fig. S5). Within the Xenium panel, 418 ligands (53.2% of distinct ligands) and 486 receptors (67.5% of distinct receptors) were identified, and we subsequently evaluated the extent of low-coverage genes among these. A Voronoi diagram visualizes the overlap of the top 25th percentile low-depth genes across cohorts, indicating that of the 418 ligands and 486 receptors, 89 (21.3%) and 80 (16.5%), respectively, were classified as lowcoverage genes (Fig. 2a). This translates to only 302 (9.3%) out of 3,234 ligand/receptor pairs being exhaustively examined using the Prime 5k. Additionally, we generated further Voronoi diagrams to illustrate the percentile rank of ligand/receptor genes across studies, thereby clarifying tissue variability (Supplementary Fig. S6).

**Figure 2.**
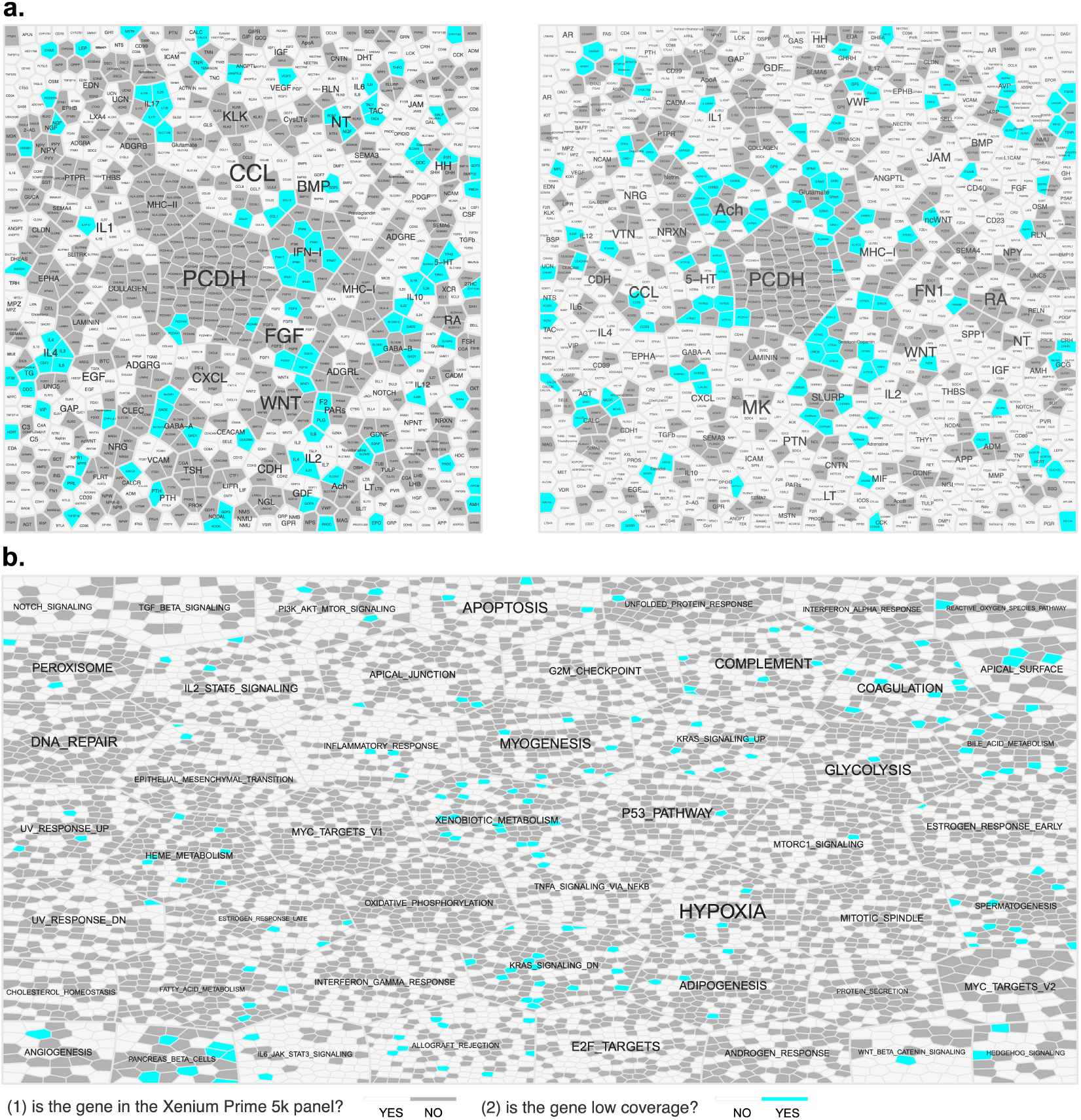
Voronoi diagrams illustrating the presence of genes on Xenium Prime 5k (white: presence; grey: absence) and their coverage (white: abundant; cyan: low-coverage) in FFPE samples. **(a)** Ligand (left) and receptor genes (right), and **(b)** Hallmark genes from the MSigDB database. Gene symbols are not shown due to the limited space but are present in Supplementary Fig. S8.

The Hallmark gene-set consists of curated gene signatures that define distinct biological processes or pathways. Among these signatures, the MYC_TARGETS _V1 exhibited the highest coverage, while the PANCREAS _BETA_CELLS genes demonstrated the lowest coverage across Xenium cohorts (Supplementary Fig. S7). Specifically, 79 genes from the MYC_TARGETS_V1 panel showed high coverage, whereas 7 (38.9%) out of 18 PANCREAS_BETA_CELLS genes were identified as low-coverage genes (Fig. 2b and S8). Notably, there were no low-coverage genes within the MYC_TARGETS_V1 signature; however, of the 200 MYC_TARGETS_V1 genes present in the panel, only 79 (39.5%) were included (Supplementary Table S2). In contrast, the OXIDATIVE _PHOSPHORYLATION signature had the lowest proportion, with only 45 (22.5%) out of 200 genes in the panel. Conversely, 69 (79.3%) out of 87 IL6_JAK_STAT3_SIGNALING genes were defined in the panel, although two of these, *CSF2* and *DNTT*, were classified as low-coverage genes.

## Discussion

This study comprehensively evaluates gene representation across pre-designed ST panels, highlighting key biases and limitations in current gene panel designs. By introducing the CI as a quantitative metric, we systematically assessed the capacity of the Xenium Prime 5k and the GeoMx WTA panels to capture gene expression across multiple datasets. Our findings reveal significant disparities in gene coverage across different biological pathways and gene categories, emphasizing the importance of panel design in ensuring robust and representative transcriptomic analyses. Furthermore, it is essential for researchers to understand any potential biases in gene coverage between panels/ST platforms when undertaking and analyzing their data.

One of the most striking observations in our study is the consistently higher CI values observed for cancer-related gene lists compared to other categories. This suggests that pre-designed ST panels are optimized for oncological research, likely due to the well-characterized nature of cancer-associated genes and their frequent inclusion in transcriptomic studies. In contrast, ligand/receptor gene pairs exhibited low CI values across most studies, indicating a potential limitation in detecting genes involved in cell-cell interaction. This finding underscores the need for enhanced panel designs, i.e., customized add-on panels, incorporating a broader representation of ligand/receptor genes. Although underrepresented low-coverage genes can still be utilized in statistical analyses within a case-control framework, researchers should consider a variety of methodologies to fully leverage the biological significance of these genes. Our study also highlights the limitations of current ST panels in accurately characterizing genes from specific protein families, such as CYP, UGT, and SLC, which exhibit complex paralogous relationships. The low CI values observed for these gene families suggest that existing panels may not sufficiently capture their diversity, potentially limiting the accuracy of pharmacogenomic and metabolic studies in spatial transcriptomics.

One of the limitations of our study is the absence of a QC process, as we relied solely on raw read counts. Advanced quality assessment and normalization methods could potentially alter the CI values. Additionally, our analysis of the ST datasets was performed without sample annotations, such as tissue type, which suggests that the CI values may be influenced by tissue or sample specificity.

In conclusion, our study provides essential insights into the design and performance of pre-designed ST panels. While current panels are well-suited for cancer research, their limitations in other disease categories or pathways highlight areas for improvement. Future advancements in ST technology should prioritize the inclusion of a more diverse gene or probe repertoire, ensuring comprehensive and unbiased transcriptomic analyses across a broader range of biological processes and disease contexts.

## Materials and Methods

### Ten gene databases

Five gene-sets were downloaded from MSigDB^12^: Hallmark (H), Pathway (C2, KEGG_LEGACY), Curated Cancer Cell Atlas (C4, 3CA), Oncogenic Signature (C6), and Cell Type Signature (C8) (accessed in January 2025). From the literature and online search, we also downloaded five additional gene lists: OMIM (Online Mendelian Inheritance in Man)^13^, GAD (Genetic Association Database)^14^, Ligand and Receptor from CellChat v2^11^, ADME (Absorption, Distribution, Metabolism, and Excretion)/PharmGKB (Pharmacogenomics Knowledgebase) genes^15^, and CYP (cytochrome P450), UGT (glucuronosyltransferase), and SLC (solute carrier) family genes (Supplementary Table S3). In the downstream analysis, we discarded gene lists with less than 10 or greater than 1,000 genes.

### Two pre-designed panels of the ST datasets

We characterized two commercialized human panels, 10X Genomics Xenium Prime 5k and Nanostring GeoMx DSP WTA, and localized two sets of ST data that are publicly available (Supplementary Table S4). We downloaded seven Xenium In Situ cohorts from the 10X Genomics data portal; seven and nine GeoMx DSP cohorts were obtained from the Nanostring web and GEO (Gene Expression Omnibus), respectively. Each platform has a consensus pipeline to assess the data quality but we do not run the pipelines to preserve transcripts/genes with low coverage. We compiled ST data into a single object using the R software (version 4.4.1). Specifically, a GeoMx data object was assembled using the following R packages: *NanoStringNCTools* (v1.12.0), *GeoMxWorkflows* (v1.10.0), and *GeomxTools* (3.8.0). On the other hand, a Xenium data object was generated using the R package *HDF5Array* (v.1.32.1).

### Coverage Index (CI) as a proxy to measure the panel coverage

We aggregated the raw read or barcode counts across samples in each cohort, and genes were sorted based on the total read counts. For each gene list from the DBs, we mapped the member genes and ranked them to use the binary classifier to evaluate the coverage depth^13^. The classifier outputs the Area Under the Receiver Operating Characteristic Curve (AUC-ROC), calculated using the R package *pROC* (v1.18.5). Here, we defined AUC-ROC as the CI, where higher CI values imply greater coverage depth. To summarize the CI values per gene list, we took a mean of the CI values across cohorts and generated a Violin/Box plot using the R package *ggplot2* (v3.5.1).

### Visualization using the Voronoi diagram

Voronoi diagrams were generated using the R package *WeightedTreemaps* (v0.1.3) to represent underrepresented genes in each gene list. We used the *voronoiTreemap* function to outline the layout and generated the diagram using the *drawTreemap* function. Two gene lists—Ligand/Receptor and KEGG pathway—were used to create the diagrams to present genes that are not present in the Xenium Prime 5k and low-coverage genes. The low-coverage genes were defined as an intersection of the 25th-percentile genes across six FFPE cohorts.

## Data and Code Availability

This study’s comprehensive analysis of public data is thoroughly documented in this published article and its accompanying Supplementary Information files. To facilitate research reproducibility and replicability, a set of scripts has been made available at https://GitHub.com/UC-ASOC/Coverage. Additionally, a ShinyApp is accessible at https://ShinyApps.UCalgary.ca/Coverage, enabling users to download (supplementary) Figures and processed data, and query users’ gene lists.

## Author contributions

HS conceived the project and HS and RK wrote the manuscript.

## Competing interests

The authors declare that there are no competing interests.

## Notes

### Competing Interest Statement

The authors have declared no competing interest.

## References

1. Burgess DJ. Spatial transcriptomics coming of age. Nat Rev Genet. 2019;20(6):317.

2. Chen J, Larsson L, Swarbrick A, Lundeberg J. Spatial landscapes of cancers: insights and opportunities. Nat Rev Clin Oncol. 2024;21(9):660–660.

3. Cheng M, Jiang Y, Xu J, et al. Spatially resolved transcriptomics: a comprehensive review of their technological advances, applications, and challenges. J Genet Genomics. 2023;50(9):625–625.

4. Lopez R, Regier J, Cole MB, Jordan MI, Yosef N. Deep generative modeling for single-cell transcriptomics. Nat Methods. 2018;15(12):1053–1058.

5. Thrupp N, Sala Frigerio C, Wolfs L, et al. Single-nucleus RNA-seq is not suitable for detection of microglial activation genes in humans. Cell Rep. 2020;32(13):108189.

6. Oszwald A, Zisser L, Schachner H, et al. Full-length target sequences of GeoMx digital spatial profiling probes reveal that gene-promiscuity predicts probe sensitivity to EDTA tissue decalcification. Sci Rep. 2024;14(1):21156.

7. Finlay JB, Ireland AS, Hawgood SB, et al. Olfactory neuroblastoma mimics molecular heterogeneity and lineage trajectories of small-cell lung cancer. Cancer Cell. 2024;42(6):1086–1086.e13.

8. Shekarian T, Ritz MF, Hogan S, et al. Multidimensional analysis of matched primary and recurrent glioblastoma identifies contributors to tumor recurrence influencing time to relapse. J Neuropathol Exp Neurol. 2025;84(1):45–45.

9. Burns GW, Fu Z, Vegter EL, et al. Spatial transcriptomic analysis identifies epithelium-macrophage crosstalk in endometriotic lesions. iScience. 2025;28(2):111790

10. Danielsen M, Emmanuel T, Nielsen MM, et al. RUNX2 as a novel biomarker for early identification of patients progressing to advanced-stage mycosis fungoides. Front Oncol. 2024;14:1421443.

11. Jin S, Plikus MV, Nie Q. CellChat for systematic analysis of cell-cell communication from single-cell transcriptomics. Nat Protoc. 2025;20(1):180–180.

12. Subramanian A, Tamayo P, Mootha VK, et al. Gene set enrichment analysis: a knowledge-based approach for interpreting genome-wide expression profiles. Proc Natl Acad Sci U S A. 2005;102(43):15545–15550.

13. Petrovski S, Wang Q, Heinzen EL, Allen AS, Goldstein DB. Genic intolerance to functional variation and the interpretation of personal genomes. PLoS Genet. 2013;9(8):e1003709.

14. Becker KG, Barnes KC, Bright TJ, Wang SA. The genetic association database. Nat Genet. 2004;36(5):431–431.

15. Park Y, Seo H, Y Ryu B, Kim JH. Gene-wise variant burden and genomic characterization of nearly every gene. Pharmacogenomics. 2020;21(12):827–827.

