## Supplementary Fig. S1 for "Evaluating gene representation in spatial transcriptomics across pre-designed panels"

**Supplemental Figure S1. Coverage indices in Curated Cancer Cell Atlas.** (top) Xenium Prime 5k and (bottom) GeoMx DSP WTA.

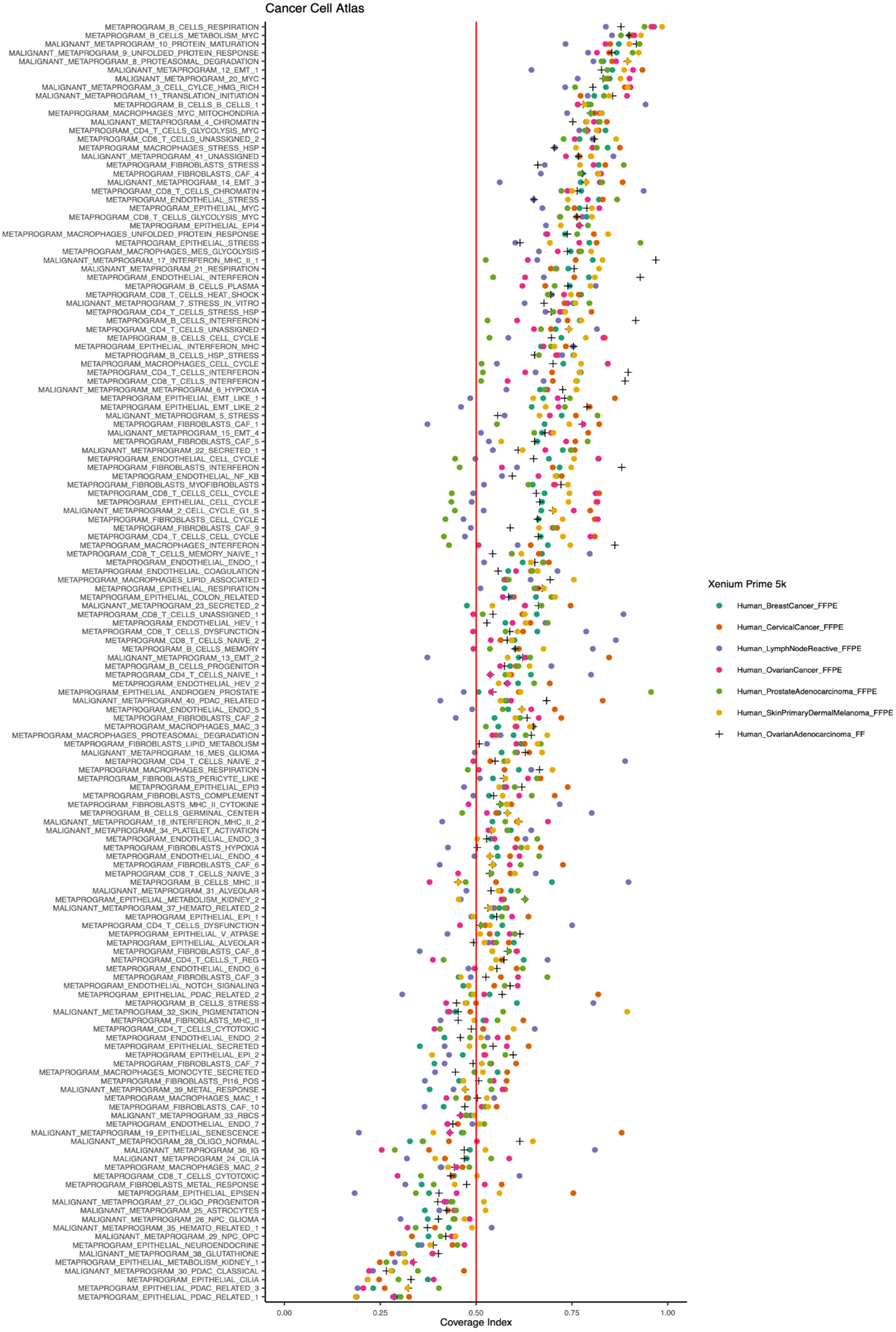

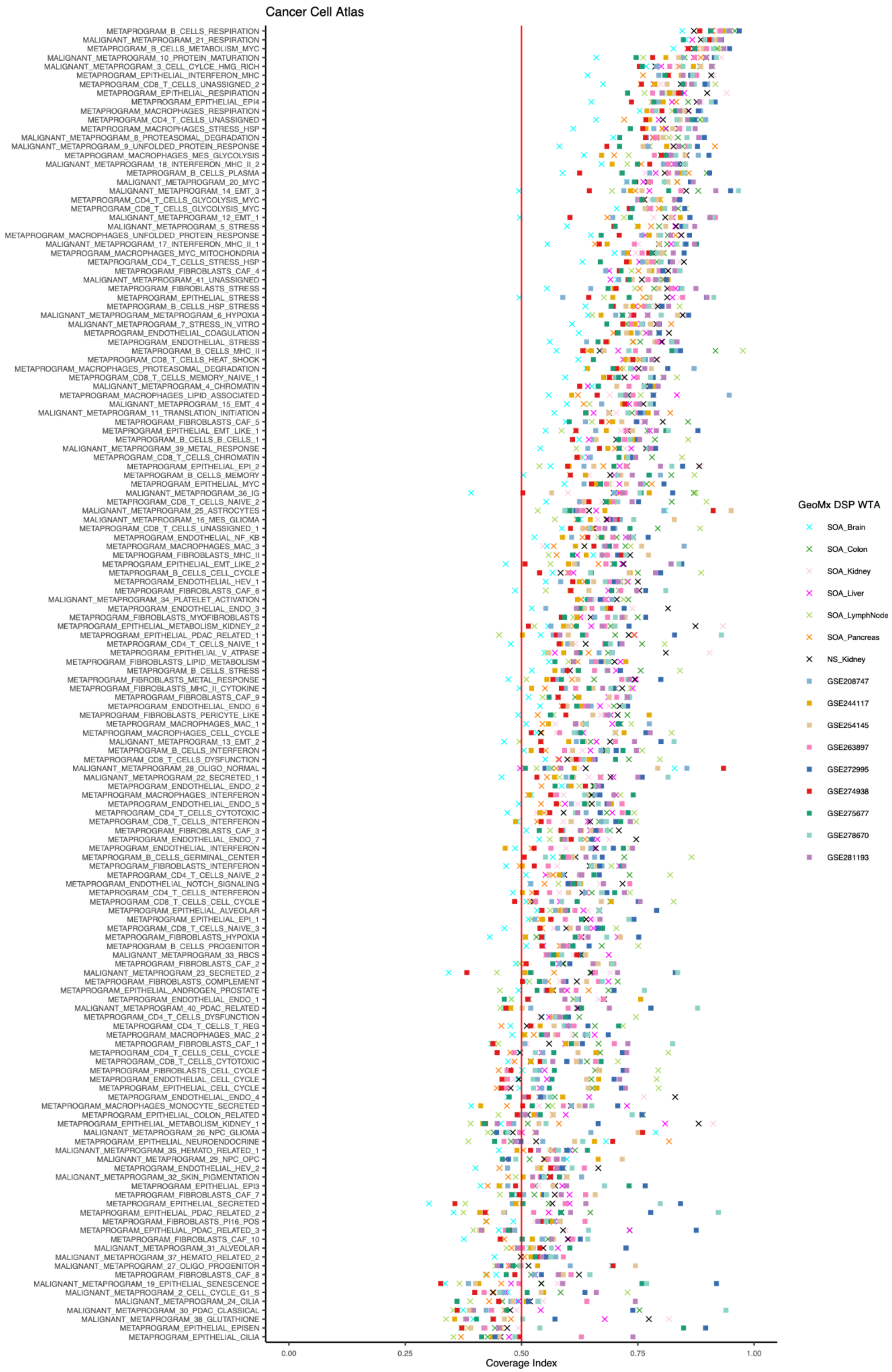
Cancer Cell Atlas
METAPROGRAM_B_CELLS_RESPIRATION
MALIGNANT_METAPROGRAM_21_RESPIRATION
METAPROGRAM_B_CELLS_METABOLISM_MYC
MALIGNANT_METAPROGRAM_10_PROTEIN_MATURATION
MALIGNANT_METAPROGRAM_3_CELL_CYLCE_HMG_RICH
METAPROGRAM_EPITHELIAL_INTERFERON_MHC
METAPROGRAM_CD8_T_CELLS_UNASSIGNED_2
METAPROGRAM_EPITHELIAL_RESPIRATION
METAPROGRAM_EPITHELIAL_EPI4
METAPROGRAM_MACROPHAGES_RESPIRATION
METAPROGRAM_CD4_T_CELLS_UNASSIGNED
METAPROGRAM_MACROPHAGES_STRESS_HSP
MALIGNANT_METAPROGRAM_8_PROTEASOMAL_DEGRADATION
MALIGNANT_METAPROGRAM_9_UNFOLDED_PROTEIN_RESPONSE
METAPROGRAM_MACROPHAGES_MES_GLYCOLYSIS
MALIGNANT_METAPROGRAM_18_INTERFERON_MHC_II_2
METAPROGRAM_B_CELLS_PLASMA
MALIGNANT_METAPROGRAM_20_MYC
MALIGNANT_METAPROGRAM_14_EMT_3
METAPROGRAM_CD4_T_CELLS_GLYCOLYSIS_MYC
METAPROGRAM_CD8_T_CELLS_GLYCOLYSIS_MYC
MALIGNANT_METAPROGRAM_12_EMT_1
MALIGNANT_METAPROGRAM_5_STRESS
METAPROGRAM_MACROPHAGES_UNFOLDED_PROTEIN_RESPONSE
MALIGNANT_METAPROGRAM_17_INTERFERON_MHC_II_1
METAPROGRAM_MACROPHAGES_MYC_MITOCHONDRIA
METAPROGRAM_CD4_T_CELLS_STRESS_HSP
METAPROGRAM_FIBROBLASTS_CAF_4
MALIGNANT_METAPROGRAM_41_UNASSIGNED
METAPROGRAM_FIBROBLASTS_STRESS
METAPROGRAM_EPITHELIAL_STRESS
METAPROGRAM_B_CELLS_HSP_STRESS
MALIGNANT_METAPROGRAM_METAPROGRAM_6_HYPOXIA
MALIGNANT_METAPROGRAM_7_STRESS_IN_VITRO
METAPROGRAM_ENDOTHELIAL_COAGULATION
METAPROGRAM_ENDOTHELIAL_STRESS
METAPROGRAM_B_CELLS_MHC_II
METAPROGRAM_CD8_T_CELLS_HEAT_SHOCK
METAPROGRAM_MACROPHAGES_PROTEASOMAL_DEGRADATION
METAPROGRAM_CD8_T_CELLS_MEMORY_NAIVE_1
MALIGNANT_METAPROGRAM_4_CHROMATIN
METAPROGRAM_MACROPHAGES_LIPID_ASSOCIATED
MALIGNANT_METAPROGRAM_15_EMT_4
MALIGNANT_METAPROGRAM_11_TRANSLATION_INITIATION
METAPROGRAM_FIBROBLASTS_CAF_5
METAPROGRAM_EPITHELIAL_EMT_LIKE_1
METAPROGRAM_B_CELLS_B_CELLS_1
MALIGNANT_METAPROGRAM_39_METAL_RESPONSE
METAPROGRAM_CD8_T_CELLS_CHROMATIN
METAPROGRAM_EPITHELIAL_EPI_2
METAPROGRAM_B_CELLS_MEMORY
METAPROGRAM_EPITHELIAL_MYC
MALIGNANT_METAPROGRAM_36_IG
METAPROGRAM_CD8_T_CELLS_NAIVE_2
MALIGNANT_METAPROGRAM_25_ASTROCYTES
MALIGNANT_METAPROGRAM_16_MES_GLIOMA
METAPROGRAM_CD8_T_CELLS_UNASSIGNED_1
METAPROGRAM_ENDOTHELIAL_NF_KB
METAPROGRAM_MACROPHAGES_MAC_3
METAPROGRAM_FIBROBLASTS_MHC_II
METAPROGRAM_EPITHELIAL_EMT_LIKE_2
METAPROGRAM_B_CELLS_CELL_CYCLE
METAPROGRAM_ENDOTHELIAL_HEV_1
METAPROGRAM_FIBROBLASTS_CAF_6
MALIGNANT_METAPROGRAM_34_PLATELET_ACTIVATION
METAPROGRAM_ENDOTHELIAL_ENDO_3
METAPROGRAM_FIBROBLASTS_MYOFIBROBLASTS
METAPROGRAM_EPITHELIAL_METABOLISM_KIDNEY_2
METAPROGRAM_EPITHELIAL_PDAC_RELATED_1
METAPROGRAM_CD4_T_CELLS_NAIVE_1
METAPROGRAM_EPITHELIAL_V_ATPASE
METAPROGRAM_FIBROBLASTS_LIPID_METABOLISM
METAPROGRAM_B_CELLS_STRESS
METAPROGRAM_FIBROBLASTS_METAL_RESPONSE
METAPROGRAM_FIBROBLASTS_MHC_II_CYTOKINE
METAPROGRAM_FIBROBLASTS_CAF_9
METAPROGRAM_ENDOTHELIAL_ENDO_6
METAPROGRAM_FIBROBLASTS_PERICYTE_LIKE
METAPROGRAM_MACROPHAGES_MAC_1
METAPROGRAM_MACROPHAGES_CELL_CYCLE
MALIGNANT_METAPROGRAM_13_EMT_2
METAPROGRAM_B_CELLS_INTERFERON
METAPROGRAM_CD8_T_CELLS_DYSFUNCTION
MALIGNANT_METAPROGRAM_28_OLIGO_NORMAL
MALIGNANT_METAPROGRAM_22_SECRETED_1
METAPROGRAM_ENDOTHELIAL_ENDO_2
METAPROGRAM_MACROPHAGES_INTERFERON
METAPROGRAM_ENDOTHELIAL_ENDO_5
METAPROGRAM_CD4_T_CELLS_CYTOTOXIC
METAPROGRAM_CD8_T_CELLS_INTERFERON
METAPROGRAM_FIBROBLASTS_CAF_3
METAPROGRAM_ENDOTHELIAL_ENDO_7
METAPROGRAM_ENDOTHELIAL_INTERFERON
METAPROGRAM_B_CELLS_GERMINAL_CENTER
METAPROGRAM_FIBROBLASTS_INTERFERON
METAPROGRAM_CD4_T_CELLS_NAIVE_2
METAPROGRAM_ENDOTHELIAL_NOTCH_SIGNALING
METAPROGRAM_CD4_T_CELLS_INTERFERON
METAPROGRAM_CD8_T_CELLS_CELL_CYCLE
METAPROGRAM_EPITHELIAL_ALVEOLAR
METAPROGRAM_EPITHELIAL_EPI_1
METAPROGRAM_CD8_T_CELLS_NAIVE_3
METAPROGRAM_FIBROBLASTS_HYPOXIA
METAPROGRAM_B_CELLS_PROGENITOR
MALIGNANT_METAPROGRAM_33_RBCS
METAPROGRAM_FIBROBLASTS_CAF_2
MALIGNANT_METAPROGRAM_23_SECRETED_2
METAPROGRAM_FIBROBLASTS_COMPLEMENT
METAPROGRAM_EPITHELIAL_ANDROGEN_PROSTATE
METAPROGRAM_ENDOTHELIAL_ENDO_1
MALIGNANT_METAPROGRAM_40_PDAC_RELATED
METAPROGRAM_CD4_T_CELLS_DYSFUNCTION
METAPROGRAM_CD4_T_CELLS_T_REG
METAPROGRAM_MACROPHAGES_MAC_2
METAPROGRAM_FIBROBLASTS_CAF_1
METAPROGRAM_CD4_T_CELLS_CELL_CYCLE
METAPROGRAM_CD8_T_CELLS_CYTOTOXIC
METAPROGRAM_FIBROBLASTS_CELL_CYCLE
METAPROGRAM_ENDOTHELIAL_CELL_CYCLE
METAPROGRAM_EPITHELIAL_CELL_CYCLE
METAPROGRAM_ENDOTHELIAL_ENDO_4
METAPROGRAM_MACROPHAGES_MONOCYTE_SECRETED
METAPROGRAM_EPITHELIAL_COLON_RELATED
METAPROGRAM_EPITHELIAL_METABOLISM_KIDNEY_1
MALIGNANT_METAPROGRAM_26_NPC_GLIOMA
METAPROGRAM_EPITHELIAL_NEUROENDOCRINE
MALIGNANT_METAPROGRAM_35_HEMATO_RELATED_1
MALIGNANT_METAPROGRAM_29_NPC_OPC
METAPROGRAM_ENDOTHELIAL_HEV_2
MALIGNANT_METAPROGRAM_32_SKIN_PIGMENTATION
METAPROGRAM_EPITHELIAL_EPI3
METAPROGRAM_FIBROBLASTS_CAF_7
METAPROGRAM_EPITHELIAL_SECRETED
METAPROGRAM_EPITHELIAL_PDAC_RELATED_2
METAPROGRAM_FIBROBLASTS_PI16_POS
METAPROGRAM_EPITHELIAL_PDAC_RELATED_3
METAPROGRAM_FIBROBLASTS_CAF_10
MALIGNANT_METAPROGRAM_31_ALVEOLAR
MALIGNANT_METAPROGRAM_37_HEMATO_RELATED_2
MALIGNANT_METAPROGRAM_27_OLIGO_PROGENITOR
METAPROGRAM_FIBROBLASTS_CAF_8
MALIGNANT_METAPROGRAM_19_EPITHELIAL_SENESCENCE
MALIGNANT_METAPROGRAM_2_CELL_CYCLE_G1_S
MALIGNANT_METAPROGRAM_24_CILIA
MALIGNANT_METAPROGRAM_30_PDAC_CLASSICAL
MALIGNANT_METAPROGRAM_38_GLUTATHIONE
METAPROGRAM_EPITHELIAL_EPISEN
METAPROGRAM_EPITHELIAL_CILIA
0.00
0.25
0.50
0.75
1.00
Coverage Index
GeoMx DSP WTA
SOA_Brain
SOA_Colon
SOA_Kidney
SOA_Liver
SOA_LymphNode
SOA_Pancreas
NS_Kidney
GSE208747
GSE244117
GSE254145
GSE263897
GSE272995
GSE274938
GSE275677
GSE278670
GSE281193
