## Supplementary Fig. S2 for "Evaluating gene representation in spatial transcriptomics across pre-designed panels"

**Supplemental Figure S2. Correlation coefficients across cohorts using the aggregated raw read counts.**

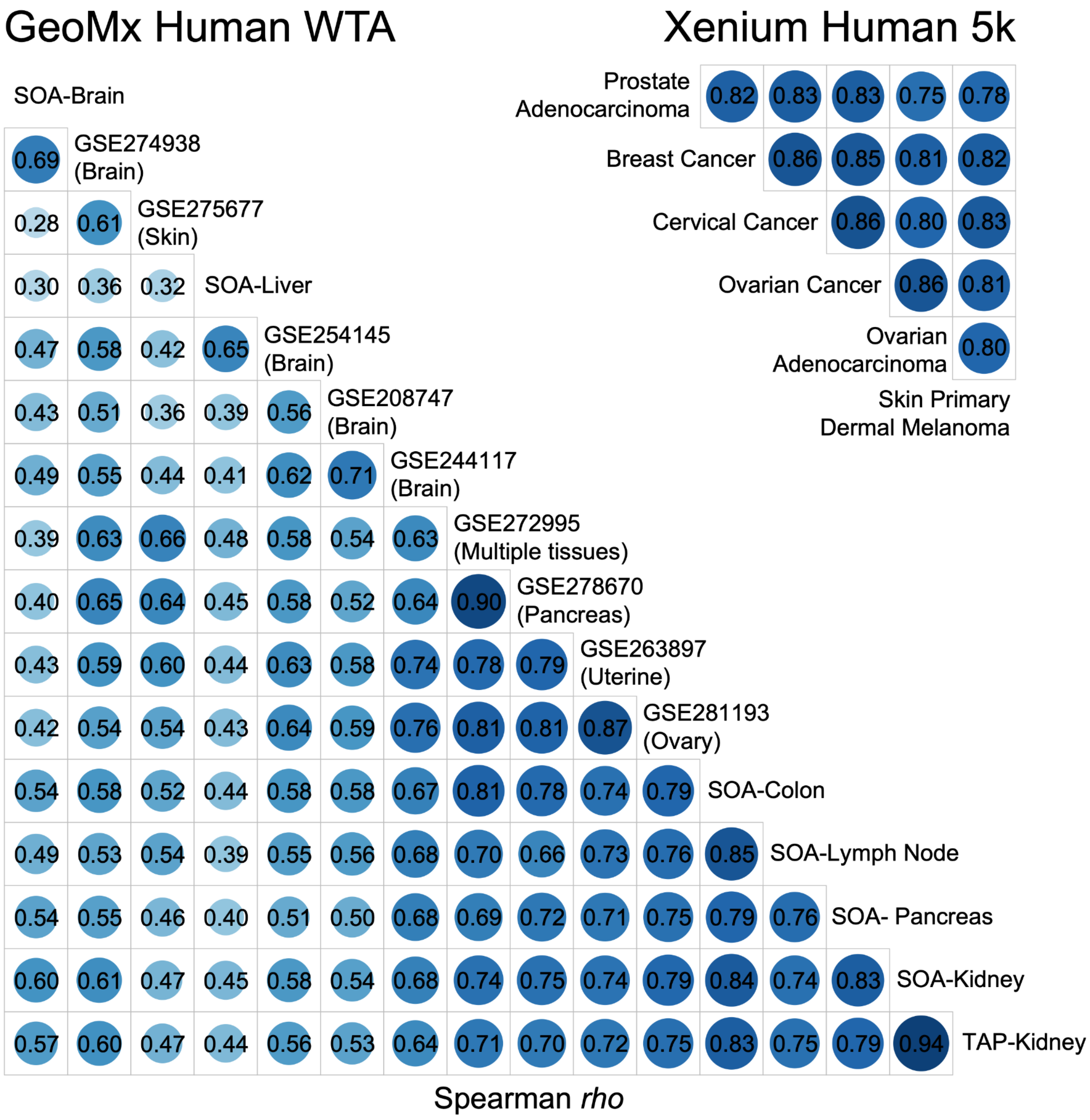
