## Supplementary figures and images for "Evaluating gene representation in spatial transcriptomics across pre-designed panels"

### Supplementary Fig. S3

**Supplemental Figure S3. Coverage indices in GAD.** (top) Xenium Prime 5k and (bottom) GeoMx DSP WTA

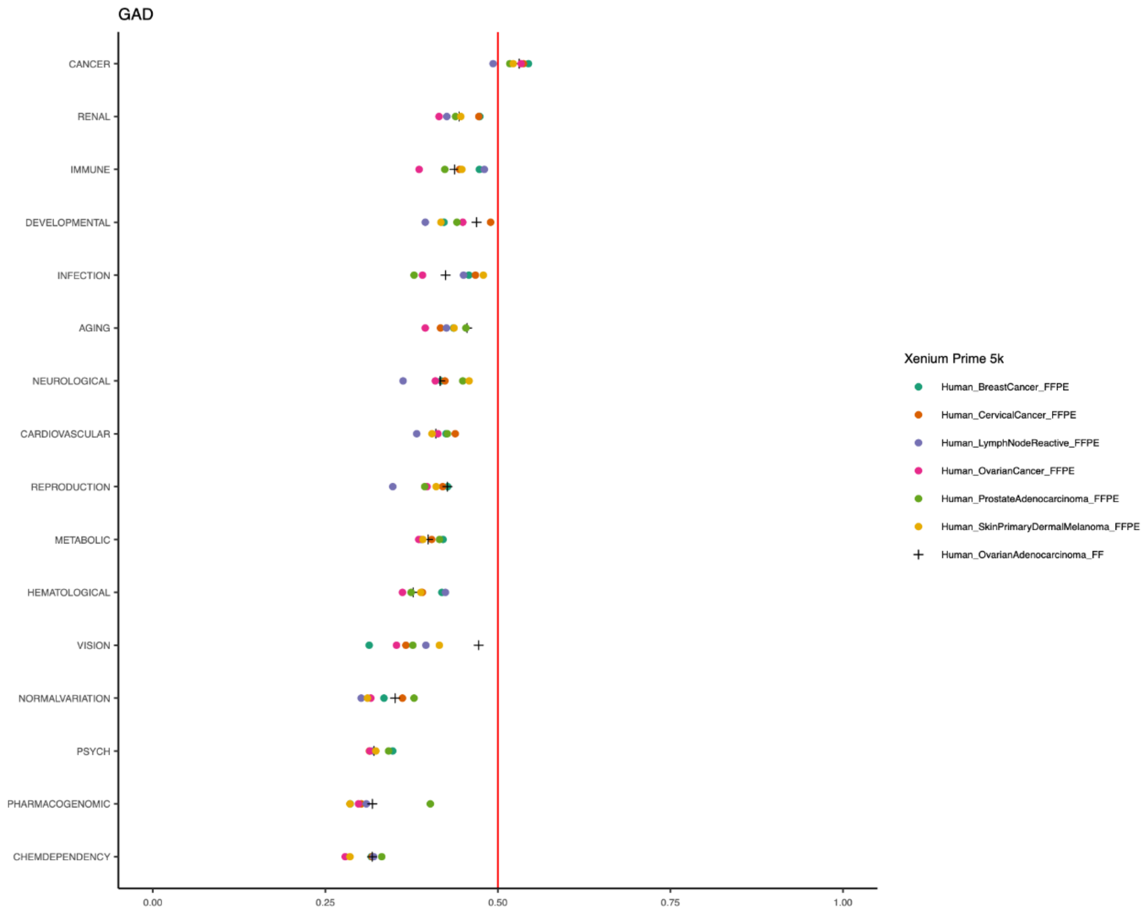


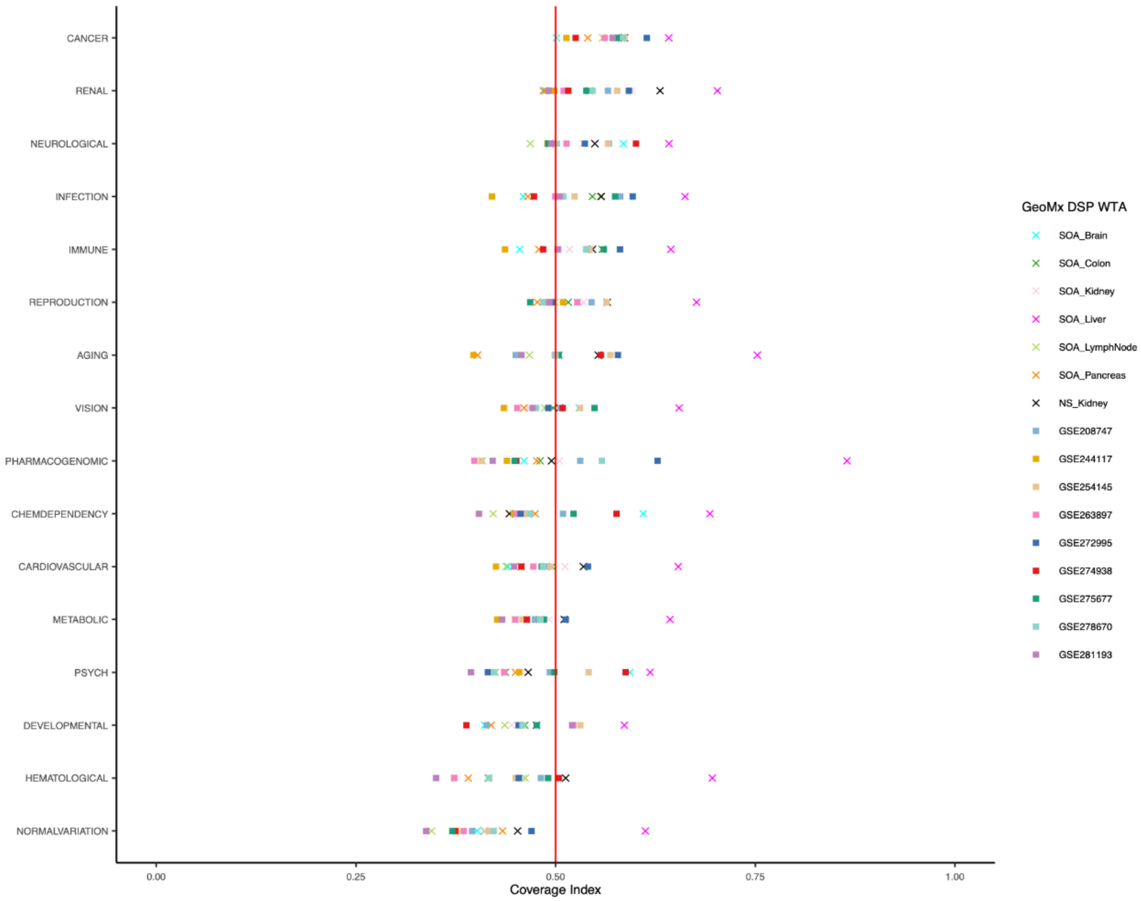

### Supplementary Fig. S4

**Supplemental Figure S4. The mean of the read counts among the six FFPE samples.**

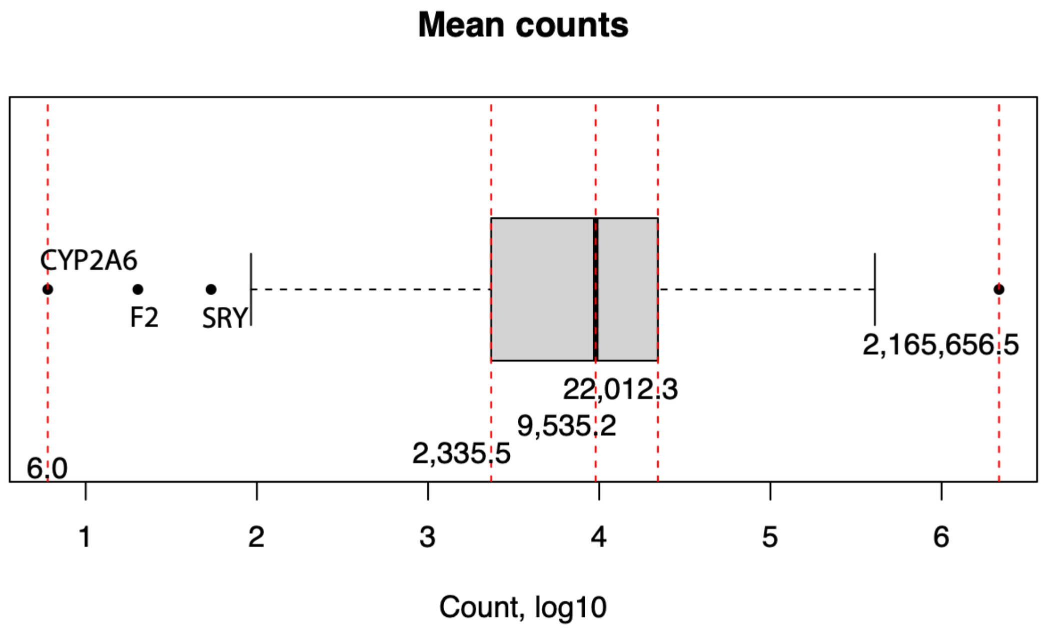

### Supplementary Fig. S5

**Supplemental Figure S5. The intersection between Ligand/Receptor genes and Xenium Prime 5k.**

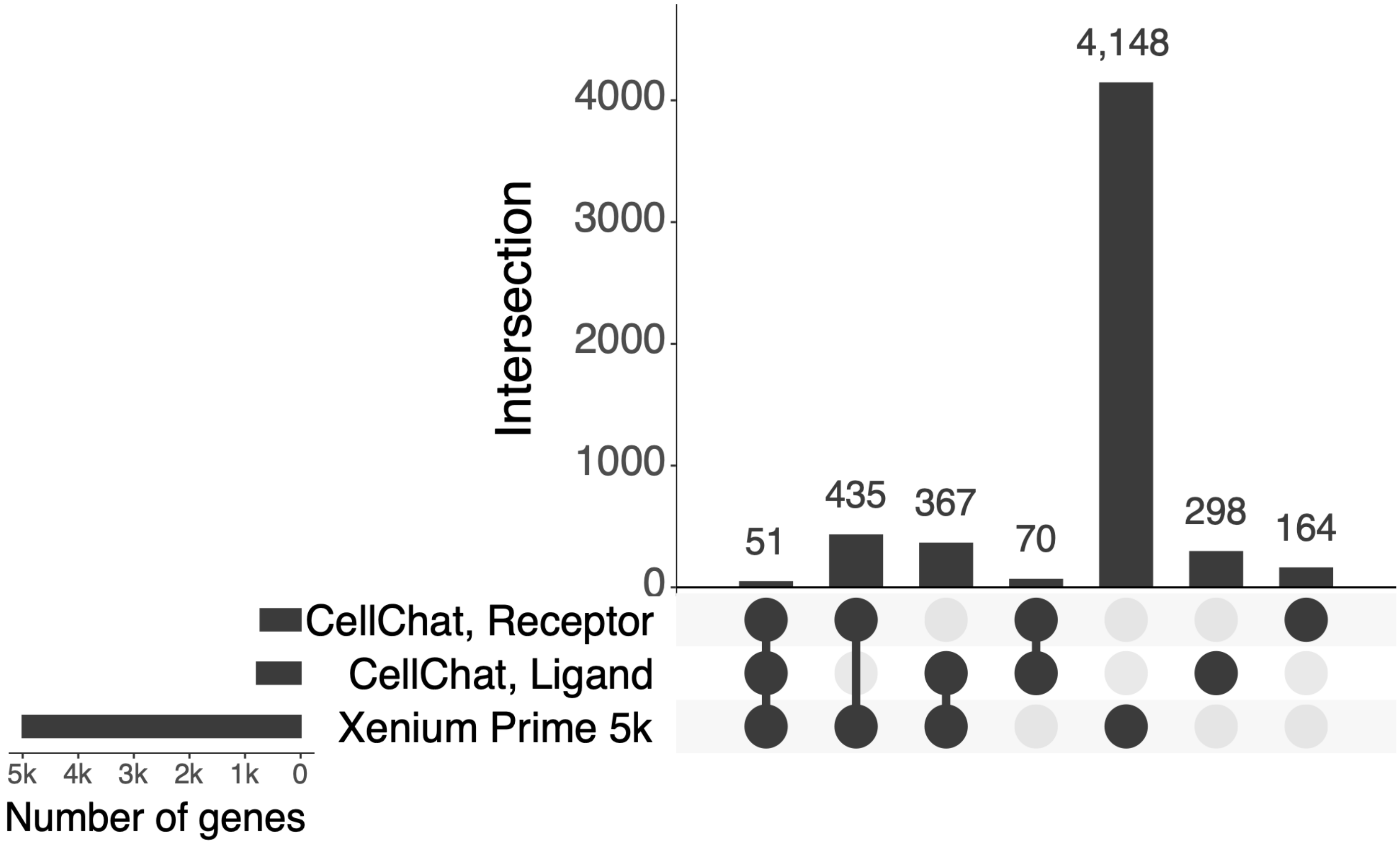

### Supplementary Fig. S7

#### Supplemental Figure S7. Coverage indices in Hallmark using the Xenium Prime 5k.

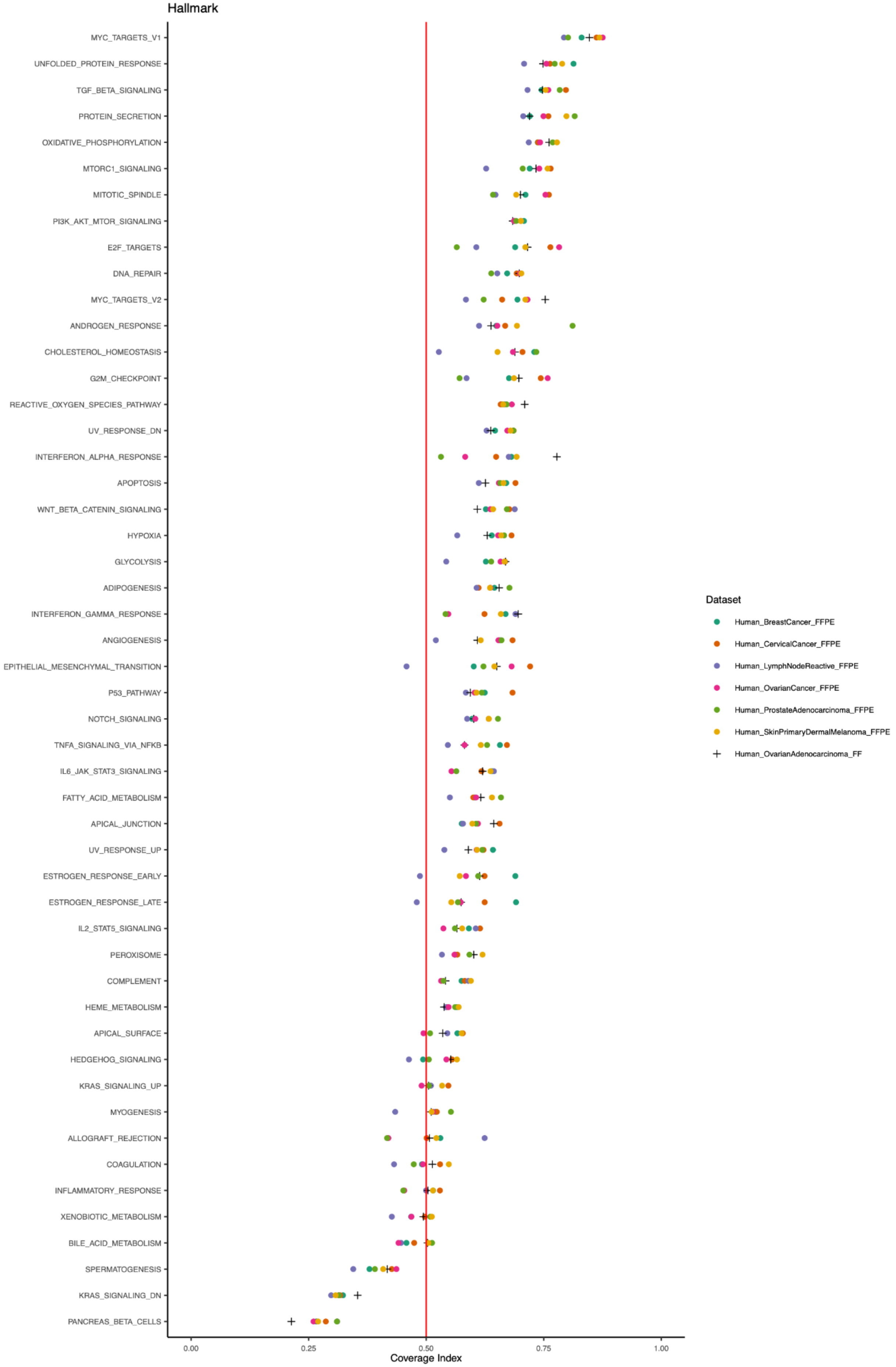
