## Supplementary Fig. S6 for "Evaluating gene representation in spatial transcriptomics across pre-designed panels"

**Supplemental Figure S6. Voronoi diagram with percentile rank in each FFPE cohort.**

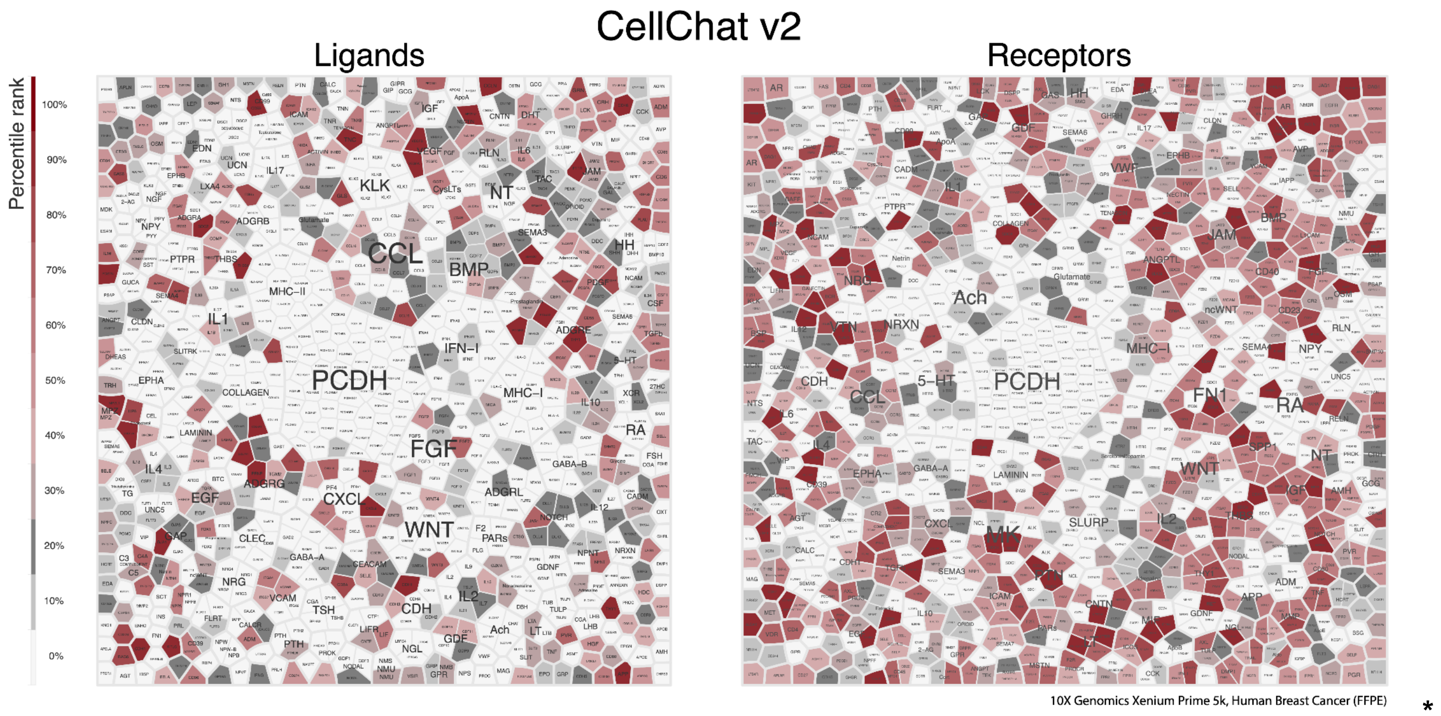


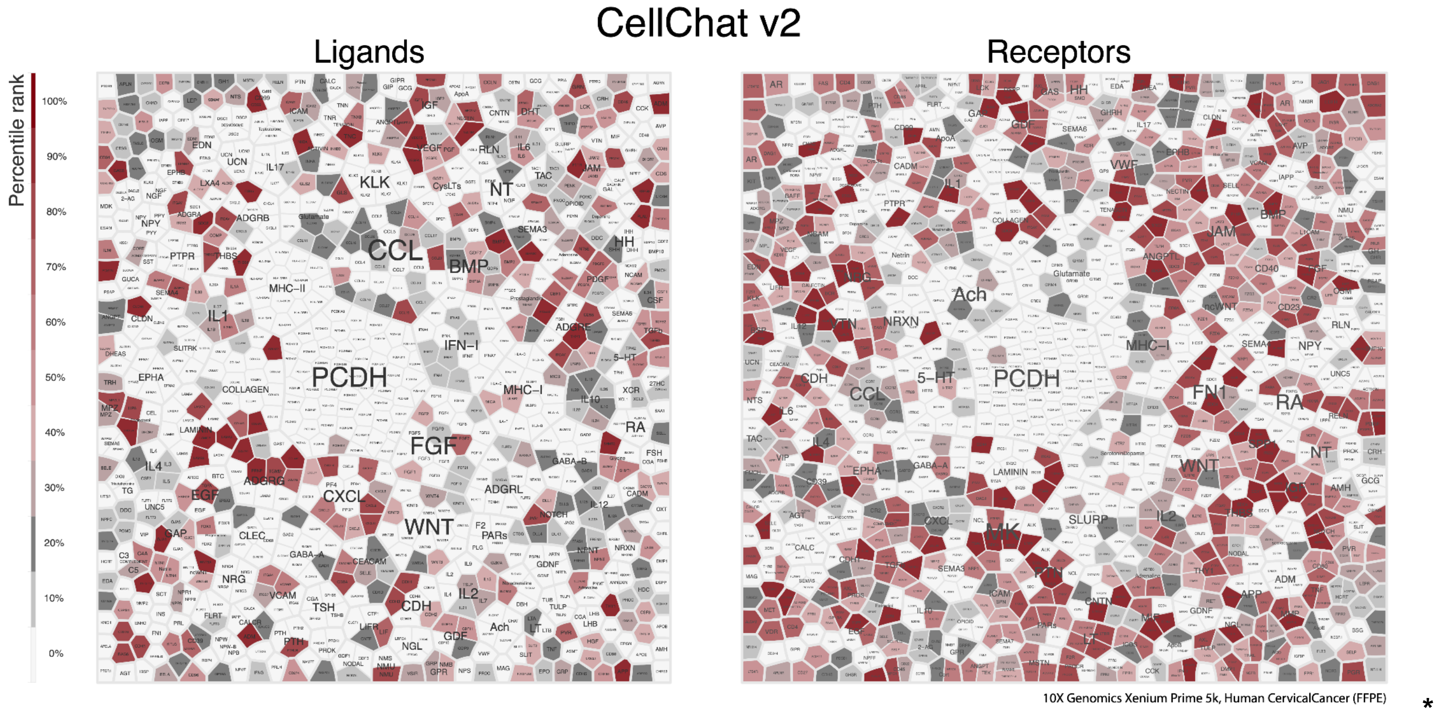

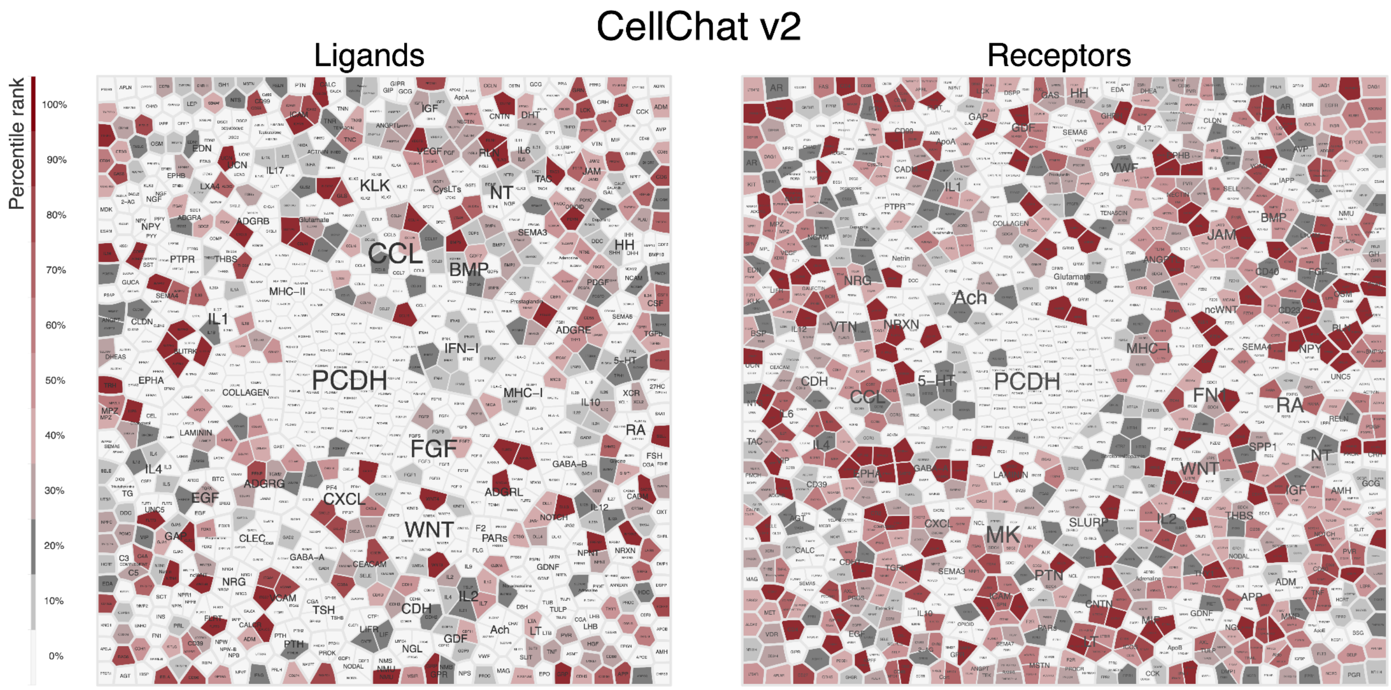


10X Genomics Xenium Prime 5k, Human Lymph Node Reactive (FFPE)

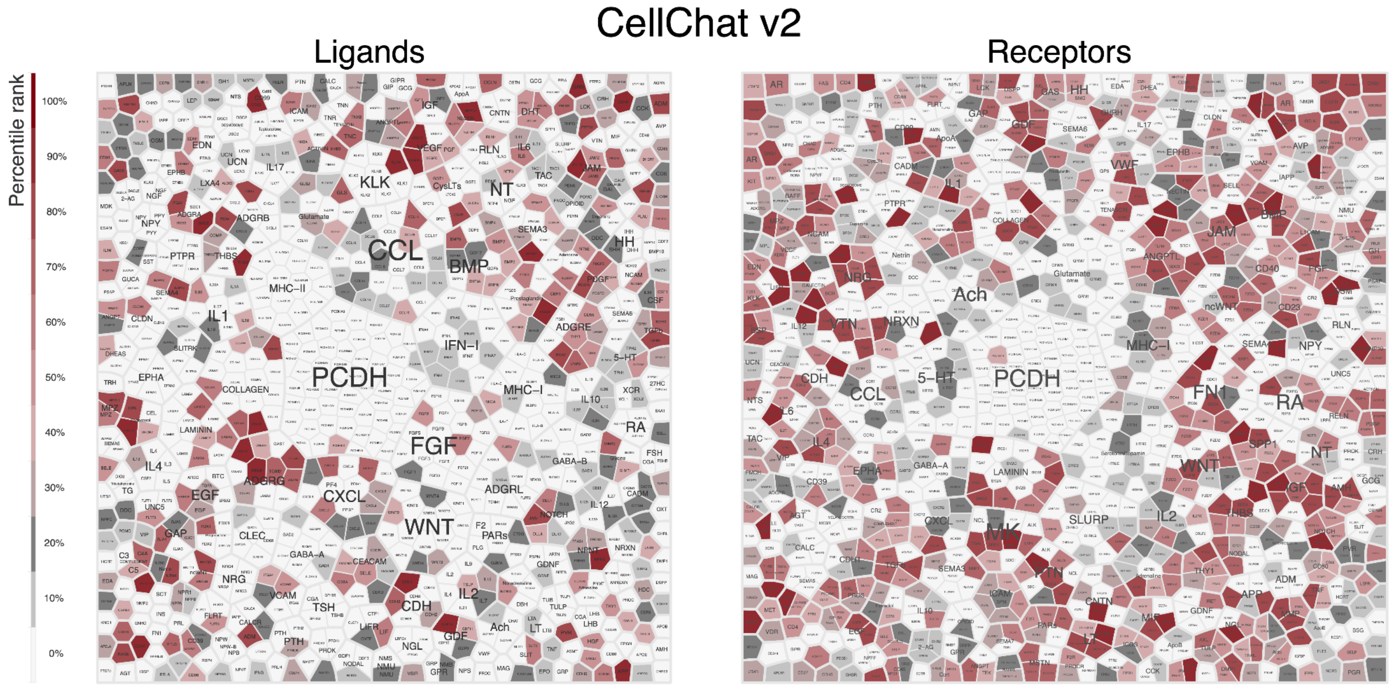


10X Genomics Xenium Prime 5k, Human Prostate Adenocarcinoma (FFPE)

### CellChat v2

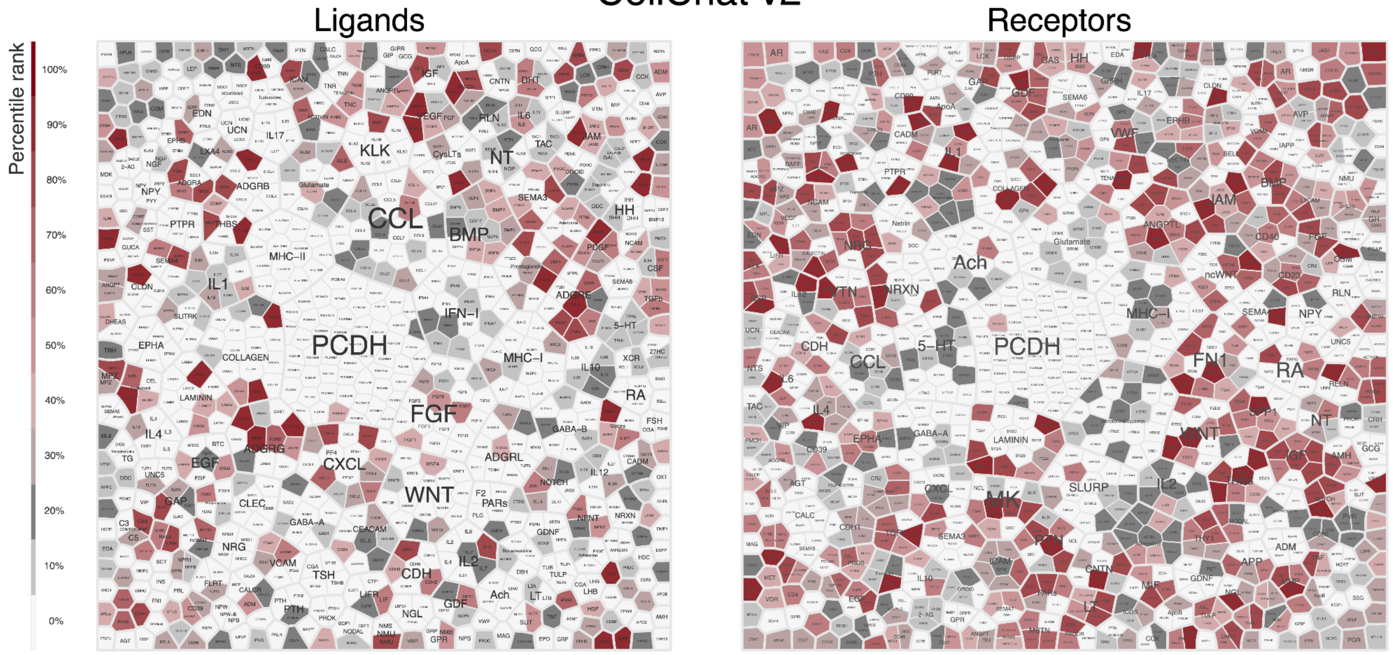


10X Genomics Xenium Prime 5k, Human Ovarian Cancer (FFPE)

### CellChat v2

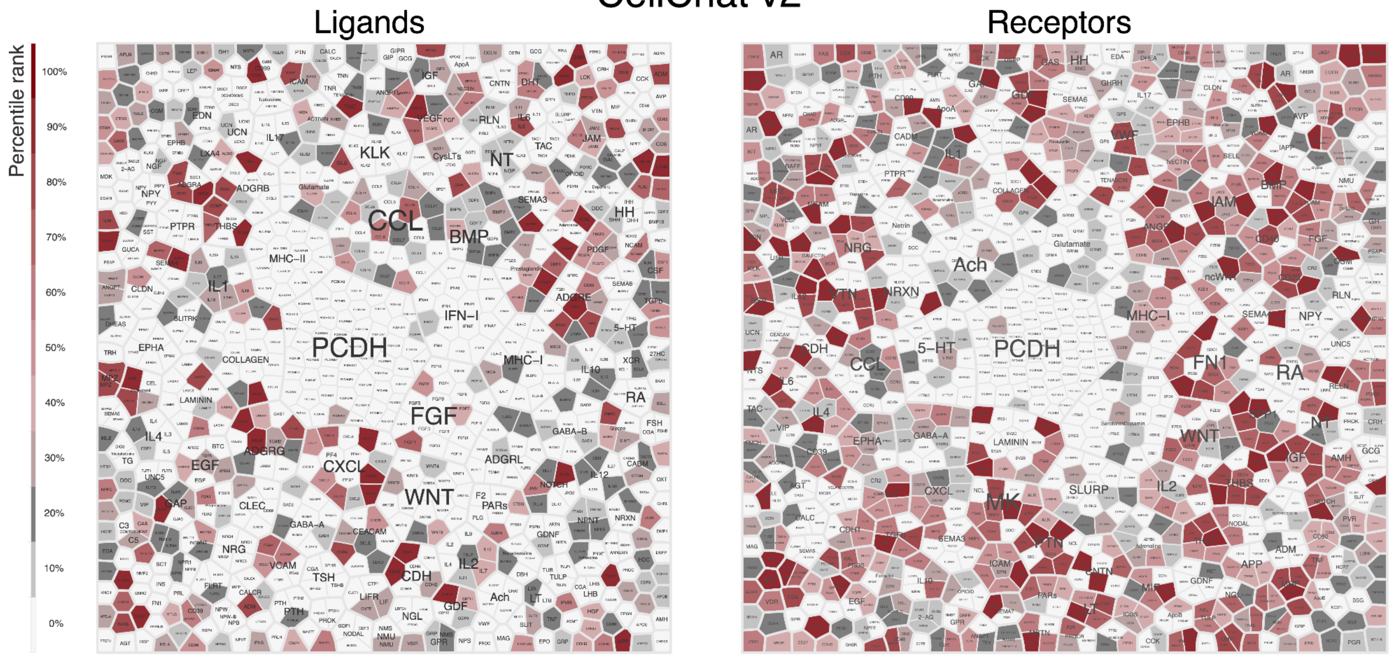


10X Genomics Xenium Prime 5k, Human Skin Primary Dermal Melanoma (FFPE)
