## Supplementary Fig. S8 for "Evaluating gene representation in spatial transcriptomics across pre-designed panels"

**Supplemental Figure S8. Voronoi diagram depicts the low-coverage genes (cyan) and Hallmark genes not presenting in the Xenium Prime 5k (grey).**

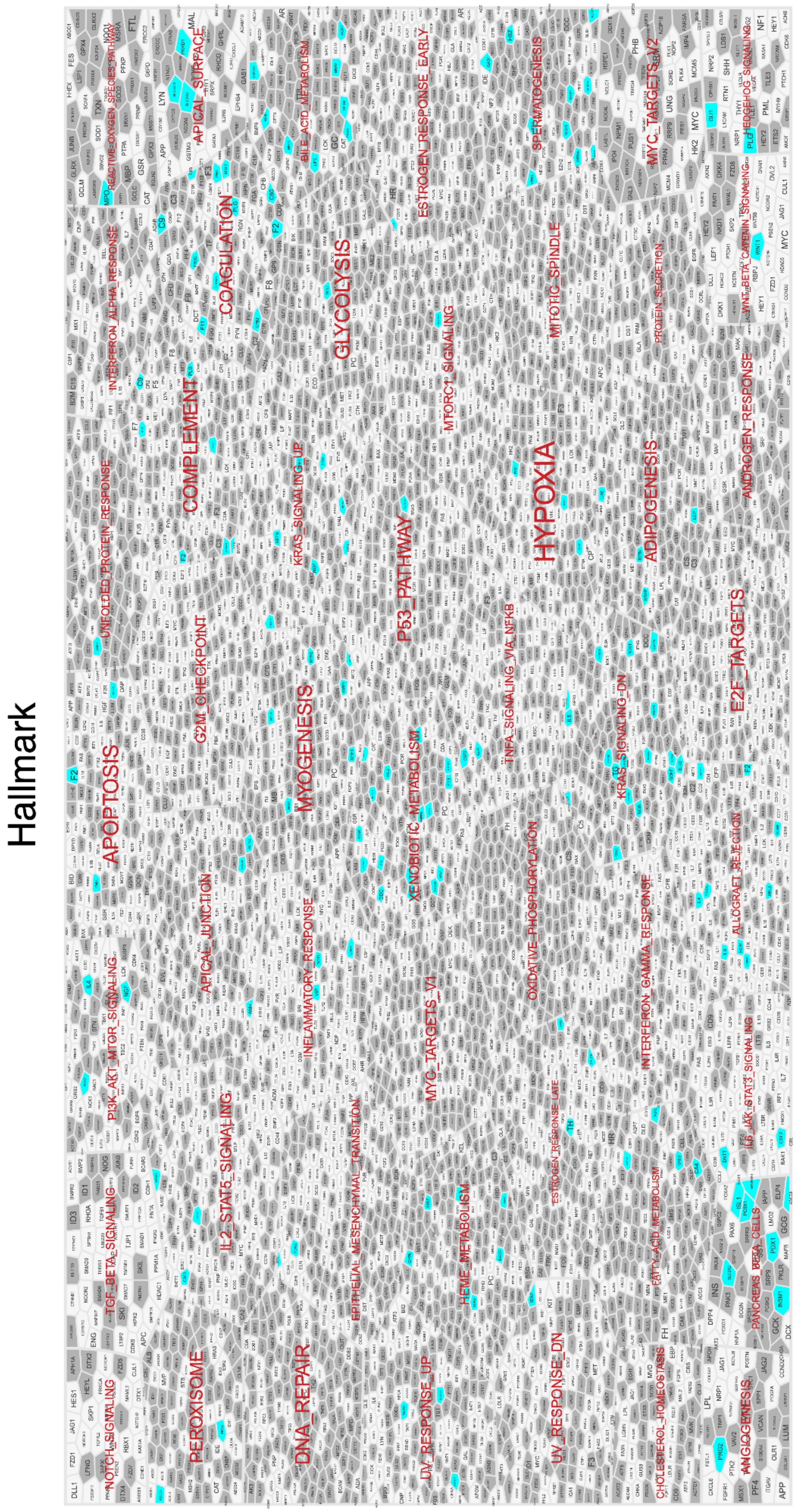
